# Activation of the TRKB receptor mediates the panicolytic-like effect of NOS inhibitor aminoguanidine

**DOI:** 10.1101/331918

**Authors:** DE Ribeiro, PC Casarotto, A Spiacci, GG Fernandes, LC Pinheiro, JE Tanus-Santos, H Zangrossi, FS Guimarães, SRL Joca, C Biojone

## Abstract

Nitric oxide (NO) triggers escape reactions in the dorsal periaqueductal gray matter (dPAG), a core structure mediating panic-associated responses, and decreases the release of BDNF *in vitro*. BDNF mediates the panicolytic effect induced by antidepressant drugs and produces these effects *per se* when injected into the dPAG. Based on these findings, we hypothesize that nitric oxide synthase (NOS) inhibitors would have panicolytic properties associated with increased BDNF signaling in the dPAG. We observed that the repeated (7 days), but not acute (1day), systemic administration of the NOS inhibitor aminoguanidine (AMG; 15 mg/kg/day) increased the latency to escape from the open arm of the elevated T-maze (ETM) and inhibited the number of jumps in hypoxia-induced escape reaction in rats, suggesting a panicolytic-like effect. Repeated, but not acute, AMG administration (15mg/kg) also decreased nitrite levels and increased TRKB phosphorylation at residues Y706/7 in the dPAG. Notwithstanding the lack of AMG effect on total BDNF levels in that structure, the microinjection of the TRK antagonist K252a into the dPAG blocked the anti-escape effect of this drug in the ETM. Taken together our data suggest that the inhibition of NO production by AMG increased the levels of pTRKB, which is required for the panicolytic-like effect observed.

## Introduction

The periaqueductal gray matter (PAG) is a midbrain structure critically associated with the expression of escape/flight responses. Chemical or electrical stimulation of dorsal regions of the PAG (dorsolateral and dorsomedial columns) in animals trigger escape reactions that have been associated with panic attacks (Depaulis et al., 1992; Brandão et al., 2008). In humans, stimulation of the dorsal PAG (dPAG) leads to a sense of fear and imminent death (Nashold et al., 1969). As evidenced by Mobbs and colleagues, situations of proximal threat increase the activation of midbrain, putatively the PAG area (Mobbs et al., 2007, 2010).

The neuronal nitric oxide synthase (NOS1 or nNOS) enzyme is highly expressed in the PAG, especially in its dorsolateral columns (Vincent and Kimura, 1992; Onstott et al., 1993). Nitric oxide (NO) can freely diffuse across cell membrane and act as retrograde signaling molecule, facilitating glutamate release in several brain regions (Lonart et al., 1992; Lawrence and Jarrott, 1993; Guevara-Guzman et al., 1994). Activation of NMDA receptor (NMDAr) by glutamate stimulates NO synthesis through calcium influx (Garthwaite et al., 1989), resulting in a positive-feedback response. Injections of NO donors or NMDA-receptor agonists into the dPAG precipitates escape reactions, while local administration of NOS inhibitors or NMDAr antagonists prevents such behaviors (De Oliveira et al., 2001; Miguel and Nunes-de-Souza, 2006; Aguiar and Guimarães, 2009).

Activation of extra-synaptic NMDAr (Vanhoutte and Bading, 2003) or increased production of NO (Canossa et al., 2002) inhibit the synthesis/release of brain-derived neurotrophic factor (BDNF). BDNF, through activation of tropomyosin-related kinase B receptors (TRKB), regulates neuronal maturation (Cohen-Cory, 2002), synaptic transmission (Figurov et al., 1996), and cell growth and survival (Ceni et al., 2014). It has been proposed that BDNF is also crucial to the effect of antidepressant drugs (Nibuya et al., 1995; Russo-Neustadt et al., 1999; Coppell et al., 2003; Duman and Monteggia, 2006; Molteni et al., 2006; Castrén et al., 2007; Martinowich and Lu, 2008). According to the ’neurotrophic hypothesis’, antidepressant drugs increase BDNF levels, especially in the hippocampus and frontal cortex, resulting in a synaptic function strengthening and, in the antidepressant/anxiolytic response (Duman et al., 1997). Antidepressant drugs such as fluoxetine are considered first line treatment for panic disorder patients (Freire et al., 2014).

Of importance to the present study, the microinjection of BDNF into the dPAG inhibited escape expression induced by electrical stimulation of this structure (Casarotto et al., 2010), or generated by the elevated T maze (ETM) (Casarotto et al., 2015), interpreted as a panicolytic-like effect (for a full description of this latter test, see (Zangrossi and Graeff, 2014). Moreover, chronic treatment with different antidepressants increased BDNF levels in the dPAG, and this effect was associated with the anti-escape action of these drugs (Casarotto et al., 2015).

Based on these findings, we here hypothesized that the inhibition of NOS would induce panicolytic-like effect by increasing BDNF signaling in the dPAG. To test this hypothesis, we first investigated whether treatment with the NOS inhibitor, aminoguanidine (AMG), inhibits escape expression in the ETM or in the hypoxia test. Next, we evaluated whether systemic AMG could decrease NO and increase BDNF and TRKB phosphorylation in the dPAG. Finally, we checked if TRKB receptors activation in dPAG was necessary for the effect of AMG systemically delivered.

## Experimental procedure

### Animals

Male Wistar rats weighing 220–250g, housed in groups of 6 were kept on a 12h dark/light cycle (lights on at 07:00 h) at 22±1°C, and given free access to food and water throughout the experiment. Independent groups of animals were used in each test. The experiments reported in this article were performed are in compliance with Brazilian Council for Animal Experimentation (CONCEA) which are based on the US National Institutes of Health guide for care and use of laboratory animals (protocol number: 114/2007 and 192/2015).

### Drugs

Aminoguanidine hydrochloride (AMG; Sigma-Aldrich, #396494, São Paulo, Brazil) was dissolved in sterile saline and injected intraperitoneally (ip) at 1ml/kg volume. K252a (Sigma-Aldrich, #05288, São Paulo, Brazil) was dissolved in 0.2% DMSO in sterile saline and microinjected into dPAG at 10pmol/200nl volume.

### Apparatus and Procedure

#### Elevated T-maze

The elevated T-maze (ETM) consisted of three arms of equal dimensions (50cm × 12cm) elevated 50cm above the floor (Zangrossi and Graeff, 2014). One arm was enclosed by 40cm high walls, perpendicular to two opposite open arms surrounded by a 1cm high Plexiglas rim.

Two days before the test the animals were gently handled by the experimenter for 5 min for habituation. Twenty-four hours before the test the rats were exposed to one of the open arms of the ETM for 30 min, as previously described (Casarotto et al., 2015). A wood barrier was temporary placed at the intersection between the open and closed arms to prevent the rat from escaping during the pre-exposition session. This pre-exposition, by shortening the latencies to withdrawal from the open arm during the test, renders the escape task more sensitive to the effects of panicolytic drugs.

The animals were tested in the elevated T-maze 60 min after the last drug or vehicle administration. For the avoidance task, each animal was placed at the distal end of the enclosed arm of the elevated T-maze facing the intersection of the arms. The time taken by the rat to leave this arm with the four paws was recorded (baseline). The same measurement was repeated in two subsequent trials (avoidance 1 and 2) at 30s intervals; during which animals were placed in a Plexiglas cage where they had been previously habituated. For the escape task (30s after avoidance 3), rats were placed at the end of the previously pre-exposed open arm and the latency to leave this arm with the four paws was recorded 3 consecutive times (escape 1, 2 and 3) with 30s intertrial intervals. A cut-off time of 300s was established for the avoidance and escape latencies. For further details of this test, see (Zangrossi and Graeff, 2014).

#### Open field

The open-field test was used to assess whether the drugs used affected the locomotion of tested animals. The test was performed in a Plexiglas circular arena (80cm diameter), with 40cm high walls. Independent groups of animals were placed in the center of the circular arena 1h after the injection for behavioral analysis during 15min. The total distance traveled (m) was determined by the ANY-MAZE software (Stoelting, USA).

#### Hypoxia test

The hypoxia chamber was a Plexiglas, roof-sealed cylinder (25cm diameter and 35cm height) with a removable rubber floor. A flow valve connected to both an air pump and a nitrogen (N_2_) cylinder was attached to the chamber, and hypoxia (7% O_2_) was produced by N_2_ administration at a flow rate of 4.5l/min during approximately 270s. The concentrations of both O_2_ and CO_2_ inside the chamber were monitored throughout the experiment (ML206 Gas Analyzer, AdInstruments, Bella Vista, NSW, Australia) and scanned online with the Power Lab Chart 5 software (AdInstruments, Bella Vista, NSW, Australia).

The animals were acclimated in the chamber under normoxic condition (21% O_2_) for 5 min. For this purpose, room air was flushed into the chamber at 4.5l/min flow rate. For the induction of hypoxia, N_2_ was flushed into the chamber (4.5l/min) for approximately 4min, until O_2_ is reduced to 7%, and maintained for 6min. The animal behavior was recorded throughout the experiment by a video camera. The number of upward jumps during the hypoxic challenge were computed by video analysis. This behavior, interpreted as escape attempts from the experimental environment, was used as a panic index (for more details of this test, see (Spiacci et al., 2015).

#### Stereotaxic surgery and local drug injection

The animals tested in experiment 5 were anesthetized with 2,2,2-tribromoethanol (250mg/kg, ip) and placed in a stereotaxic frame. A guide cannula made of stainless steel (OD 0.6mm) was implanted into the brain aimed at the dPAG. The following coordinates from lambda were used (Paxinos and Watson, 2007): AP = 0mm, ML = +1.9mm, DV = −3.2mm. The guide cannula was fixed to the skull with acrylic resin and two stainless steel screws. A stylet with the same length as the guide cannula was fixed in the resin to prevent obstruction.

At the end of the surgery, all animals received an antibiotic preparation (0.3ml, intramuscular, of benzylpenicillin and streptomycin, Pentabiotico Veterinário Pequeno Porte; Forte Dodge, Brazil), and flunixin meglumine (0.1ml subcutaneous, Banamine, Schering–Plough, Brazil) for post-surgery analgesia. The animals were left undisturbed for 5 days after the surgery, except for normal handling for cage cleaning.

For drug microinjection, a dental needle (0.3mm outer diameter) was introduced through the guide cannula until its tip was 1mm below the cannula end. A volume of 200nl was injected into the DPAG for 60s using a 10ul microsyringe (Hamilton, USA) attached to a microinfusion pump (KD Scientific, USA). The displacement of an air bubble inside the polyethylene catheters connecting the syringe to the intracerebral needles was used to monitor the microinjection. The needle was removed 30s after the injection was finished. In order to confirm the site of injection, after the end of the experiment all animals were deeply anesthetized with 4% chloral hydrate (1ml/100g, ip) and 200nl of fast green dye was injected into the PAG. The brains were removed and sliced for determination of the injection site. Only animals with needle tips inside the dorsolateral PAG were considered for the experimental analysis.

### Sample collection and preparation

#### BDNF assay and western-blotting protocol

Independent groups of animals were deeply anesthetized with chloral hydrate (4%) and the dorsal periaqueductal gray matter (dPAG) dissected using punching needles (OD 4mm). The samples were homogenized in lysis buffer (137mM NaCl; 20mM Tris-HCl; 10% glycerol) containing protease inhibitor cocktail (Sigma Aldrich, USA, #P2714) and sodium orthovanadate (0.05%, Sigma Aldrich, USA, S6508). The homogenate was centrifuged (10000 × g) at 4°C for 15min and the supernatant collected and stored at −80°C.

The BDNF levels in the PAG samples were assessed by commercial sandwich ELISA kit (Promega, #G7610, USA) according to the manufacturer’s instructions. Briefly, following preincubation with monoclonal anti-BDNF antibody and blockade of non-specific sites, the samples were added to 96-well plates. After the secondary and tertiary antibodies, the developed color intensity was read at 450nm and compared to a standard recombinant BDNF curve (7.8-500pg/ml). The BDNF concentration was normalized by total protein content in each sample.

The levels of TRKB and phosphorylated TRKB (pTRKB) were determined by western blotting (Saarelainen et al., 2003). Briefly, thirty micrograms of total protein content in each sample was separated in acrylamide gel electrophoresis and transferred to a PVDF membrane. Following blockade with 5%BSA in TBST (20mM Tris-HCl; 150mM NaCl; 0.1% Tween20) the membranes were exposed to anti-pTRK at Tyr706/707 residues (1:1000, Cell Signaling, #4621, USA), total TRKB (1:1000, Cell Signaling, #4603, USA) or GAPDH (1:2000, Santa Cruz, USA, #sc25778) overnight at 4 °C. After washing with TBST the membranes were incubated with secondary HRP-conjugated anti-rabbit IgG (1:2000, Santa Cruz, #sc2317, USA). Following washing with TBST and TBS the membranes were incubated with 4-chloronaphtol (4CN, Perkin Elmer, #NEL300001EA, USA) for color development. The dried membranes were scanned and the intensity of bands relative to pTRKB, TRKB and GAPDH were determined using ImageJ software (NIH, version 1.47, USA).

#### Quantification of nitrite concentration in PAG

Samples from dPAG were homogenized in ice-cold phosphate buffer (pH 7.4; 0.5ml) and kept on dark until use (within 5 minutes). Nitrite concentrations were analyzed in duplicate through an ozone-based reductive chemiluminescence assay as previously described (Feelisch et al., 2002). Briefly, 400ul of dPAG samples were injected in a purge vessel, and approximately 8ml triiodide solution (2g potassium iodide and 1.3g iodine dissolved in 40ml water with 140ml acetic acid) was added. The triiodide solution reduces nitrites to NO, which is purged by ozone gas and detected by NO analyzer (Sievers Model 280 NO analyzer, Boulder, CO, USA). The data was analyzed using the software OriginLab (version 6.1).

### Experimental Design

#### Experiment 1: effect of AMG acute injection in ETM

At day 1, independent groups of experimentally naive rats were pre-exposed to one of the open arms as described previously. On the 2nd day, the animals received a single administration of AMG (15, 30 or 60m/kg, ip) or vehicle solution and were tested in the ETM 1 hour after the drug injection. This dose range was based on previous studies from our group (Montezuma et al., 2012).

#### Experiment 2: effect of AMG repeated treatment in the ETM, hypoxia-induced escape, and open-field test (OF)

Independent groups of experimentally naive rats were used for each behavioral test. AMG (15, 30 or 60m/kg) or vehicle solution was injected ip once a day for 7 days. All the behavioral analysis were conducted 1h after the last injection of AMG or vehicle solution.

#### Experiment 3: effect of AMG treatment on BDNF levels in the PAG

Independent groups of experimentally naive animals received daily ip injections of AMG (15, 30 or 60mg/kg) or vehicle solution for 1 or 7 days. dPAG samples were collected as described previously 1h after the last injection.

#### Experiment 4: effect of AMG treatment on pTRKB and nitrite levels in the PAG

Independent groups of experimentally naive rats received daily ip injections of AMG (15mg/kg) or vehicle solution for 1 or 7 days. dPAG samples were collected as described above 1h after the last injection. Since no changes were observed in the levels of GAPDH or total TRKB (data not shown), the intensity relative to pTRKB were normalized by total TRKB and expressed as percent from control group.

A separate cohort of animals received daily ip injections of AMG (15mg/kg) or vehicle solution for 7 days. On day 6, half of vehicle-treated animals received AMG, therefore the following groups were: vehicle, AMG-acute, AMG-repeated. 1h after the last injection, the animals were euthanized for quantification of nitrite levels in the dPAG, as described above.

#### Experiment 5: effect of intra PAG K252a on AMG-induced panicolytic-like effect

To test the possible causal relationship between AMG-induced behavioral effects and the changes in pTRKB levels in the dPAG, experimentally naïve animals with a surgically implanted cannula aiming to dPAG were treated ip with AMG (15 mg/kg) or saline for 7 days and submitted to the ETM 1h after the last injection. On days 1, 4 and 7 the animals received intra-dPAG injections of K252a (10 pmol/200nl) or vehicle solution 10 min before the ip administration of AMG. The final groups were: vehicle-saline; vehicle-AMG; K252a-saline; K252a-AMG.

### Statistical Analysis

Data obtained in ETM was analyzed by two-way ANOVA, with drug treatments and trials as factors, followed by Newman-Keuls’ post test, when appropriate. Specifically for experiment 5, data was analysed by multivariate ANOVA (factors: trials, systemic treatment, intra-dPAG injection) followed by Newman-Keuls’ post test. Results obtained in the open field and ELISA were analyzed by one-way ANOVA, followed by Newman-Keuls’ post test. Nitrite levels in dPAG and hypoxia-induced escape reactions were analyzed by Kruskal-Wallis test, followed by Dunn’s post hoc. Western blotting data was analyzed by Mann-Whitney’s test. For all experiments a p<0.05 was considered statistically significant. SPSS 25.0 and GraphPad Prism 6 softwares were used for statistical analysis and graphical representation of the data.

### Results

#### Experiment 1: effect of AMG acute injection on elevated T maze (ETM)

The two-way repeated measures ANOVA of avoidance data in acutely treated groups showed a significant effect of trials [F(2,29)= 42.45; p<0.05], but no effect of treatment [F(3,29)= 0.36; not significant – NS] or interaction between these factors [F(6,29)= 0.38; NS] (table 1). No effect of trials [F(2,29)= 0.17; NS], treatment [F(3,29)= 0.49; NS] or interaction [F(6,29)= 0.22; NS] were found for escape latencies, as seen in figure 1a.

**Table 1:**
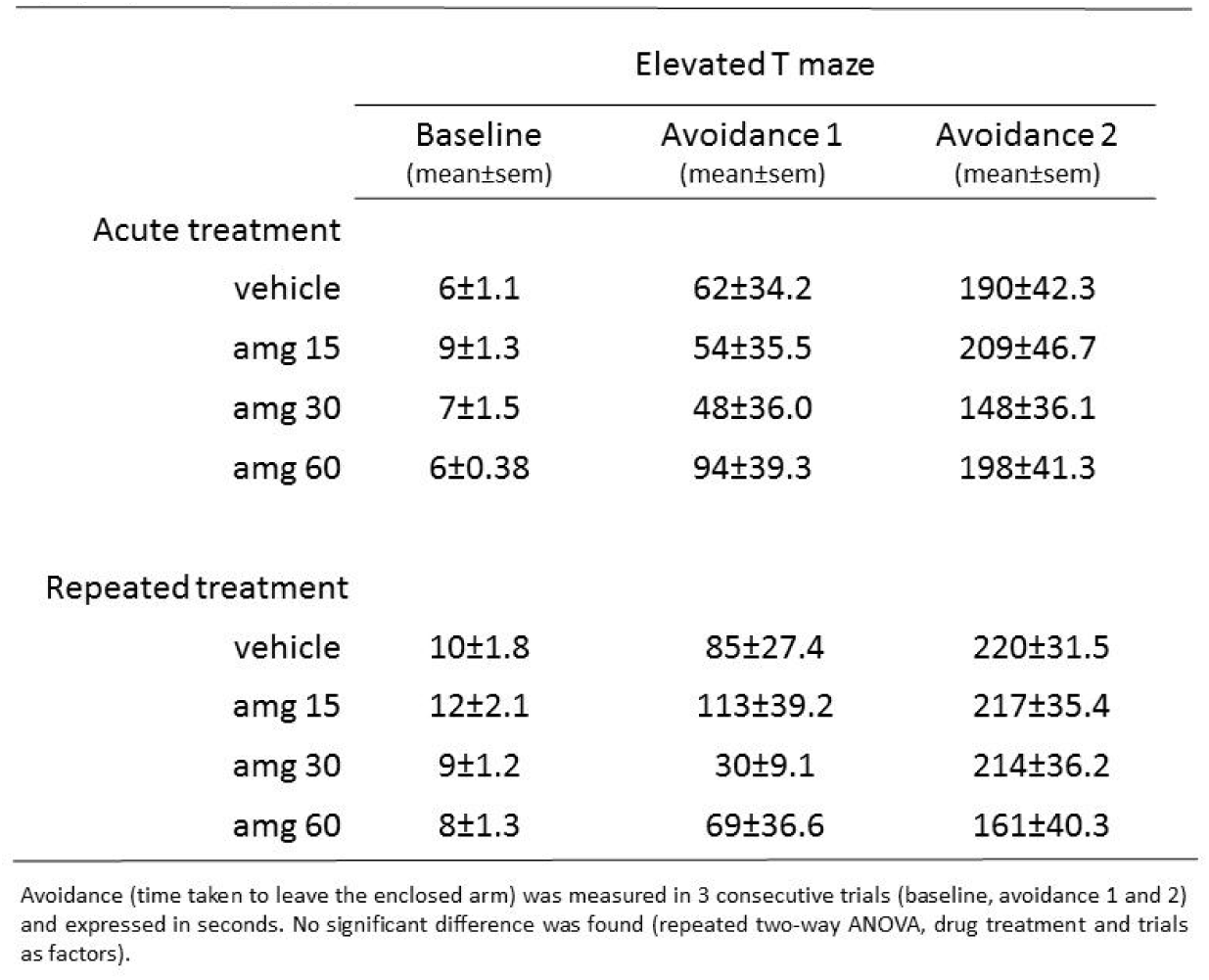
Effect of acute and repeated treatment with aminoguanidine (15,30, 60 mg/kg i.p.) on avoidance task in elevated T maze

|  |  | Elevated T maze |  |  |
| --- | --- | --- | --- | --- |
|  |  | Baseline<br>(mean±sem) | Avoidance 1<br>(mean±sem) | Avoidance 2<br>(mean±sem) |
| Acute treatment |  |  |  |  |
|  | vehicle | 6±1.1 | 62±34.2 | 190±42.3 |
|  | amg 15 | 9±1.3 | 54±35.5 | 209±46.7 |
|  | amg 30 | 7±1.5 | 48±36.0 | 148±36.1 |
|  | amg 60 | 6±0.38 | 94±39.3 | 198±41.3 |
| Repeated treatment |  |  |  |  |
|  | vehicle | 10±1.8 | 85±27.4 | 220±31.5 |
|  | amg 15 | 12±2.1 | 113±39.2 | 217±35.4 |
|  | amg 30 | 9±1.2 | 30±9.1 | 214±36.2 |
|  | amg 60 | 8±1.3 | 69±36.6 | 161±40.3 |

Avoidance (time taken to leave the enclosed arm) was measured in 3 consecutive trials (baseline, avoidance 1 and 2) and expressed in seconds. No significant difference was found (repeated two-way ANOVA, drug treatment and trials as factors).

**Figure 1.**
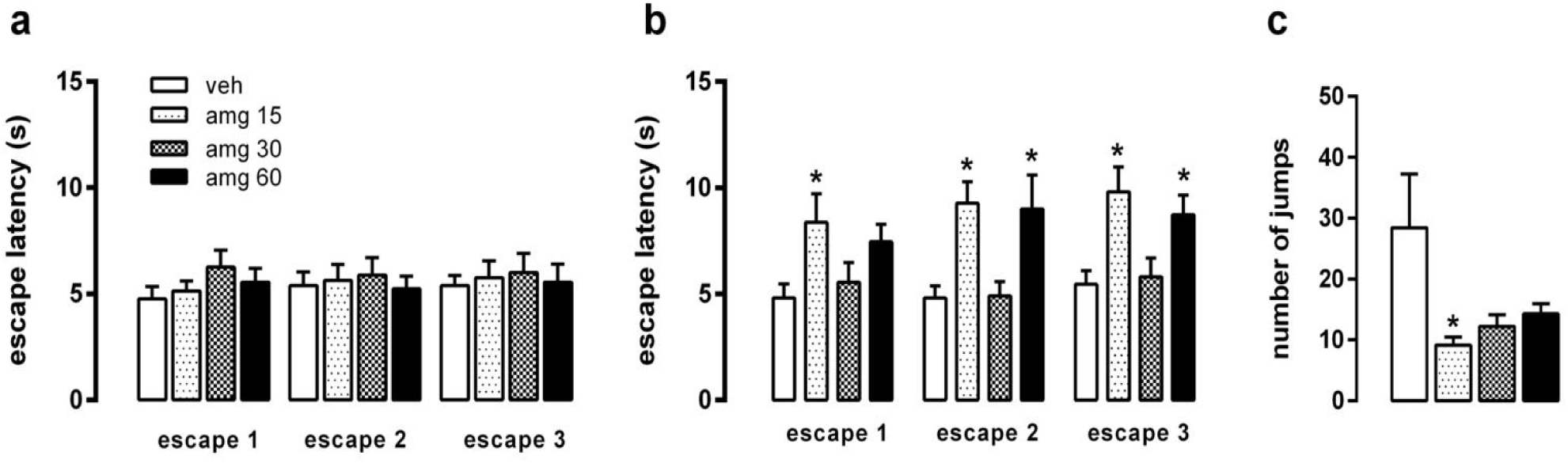
Repeated treatment with aminoguanidine (amg) induces panicolytic-like effect in the ETM and hypoxia test. Effect of amg after 1 (a: n=8,8,8,9) or 7 days treatment (b: n=11/group) on escape latency in ETM, and in the hypoxia-induced escape reaction (c: n=7,8,8,8). Columns represent mean ± SEM; two-way ANOVA (a, b) followed by Newman-Keuls’ multiple comparison test (when applicable) or Kruskal-Wallis test (c), *p^<^0.05 *vs* vehicle-treated group at the same trial.

#### Experiment 2: effect of AMG repeated treatment on the ETM, open field and hypoxia-induced escape

The analysis of avoidance data in AMG repeatedly treated groups also showed a significant effect of trials [F(2,24)= 57.94; p<0.05], but no effect of treatment [F(3,24)= 0.72; NS] or interaction between these factors [F(6,24)= 1.14; NS], as shown in table 1. On the other hand, the analysis of escape latencies data (fig. 1b) revealed a significant effect of treatment [F(3,24)= 3.63; p<0.05], without effects of trials [F(2,24)= 2.79; NS] or interaction between these factors [F(6,24)= 0.89; NS]. The post-hoc test showed that AMG at 15 and 60 mg/kg significantly increased the latency to leave the open arm (Newman-Keuls, p<0.05). For additional experiments, the lower effective dose of 15mg/kg was chosen.

AMG treatment did not induce locomotor changes in the open field test (F(3,24)= 0.07, NS]. The mean±SEM(n) of total distance traveled, in meters, were vehicle: 22.61±6.73(6); AMG 15mg: 21.09±4.63(7); AMG 30mg: 22.38±5.68(6) and AMG 60mg: 19.63±0.86(6).

The Kruskal-Wallis test showed a significant effect of AMG treatment in hypoxia test [H= 7.82; p= 0.04] (fig. 1c). AMG 15mg/kg reduced the number of jumps compared to vehicle-treated group (Dunn’s p<0.05).

#### Experiment 3: effect of AMG treatment on BDNF levels in the PAG

One-way ANOVA revealed that neither acute [F(3,19)= 0.96; NS] nor repeated treatment with AMG [F(3,39)= 0.34; NS] significantly changed BDNF levels in the dPAG, (table 2).

**Table 2:** Effect of acute and repeated treatment with aminoguanidine (15,30, 60 mg/kg i.p.) on BDNF and total TRKB levels in dPAG

|  |  | BDNF<br>(mean±sem) | TRKB<br>(mean±sem) |
| --- | --- | --- | --- |
| Acute treatment |  |  |  |
|  | vehicle | 100±14.5 | 100±12.3 |
|  | amg 15 | 103±14.5 | 134±26.2 |
|  | amg 30 | 77±8.7 | - |
|  | amg 60 | 88±10.4 | - |
| Repeated treatment |  |  |  |
|  | vehicle | 100±13.9 | 100±10.3 |
|  | amg 15 | 109±10.0 | 92.0±10.0 |
|  | amg 30 | 93±13.8 | - |
|  | amg 60 | 104±7.6 | - |

BDNF and TRKB levels are expressed as percentage of the vehicle group. TRKB levels were not assessed in amg 30 and amg 60 groups. No significant difference was found by one-way ANOVA (BDNF analysis) and t Test (TRKB analysis).

#### Experiment 4: effect of AMG treatment on pTRKB and nitrite levels in the PAG

As shown in figure 2a, Mann-Whitney’s U test indicates no significant effect of acute treatment with AMG on pTRKB/TRKB levels [U= 7.00; p= 0.18] in the dPAG. On the other hand, a significant effect on pTRKB/TRKB levels [U= 2.00; p= 0.03] was observed after repeated drug treatment (fig. 2b). Regarding nitrite levels in the dPAG, AMG decreased its levels after repeated but not acute administration (fig. 2c), as indicated by Kruskal-Wallis [H= 6.33; p= 0.04], followed by Dunn’s multiple comparison test.

**Figure 2.**
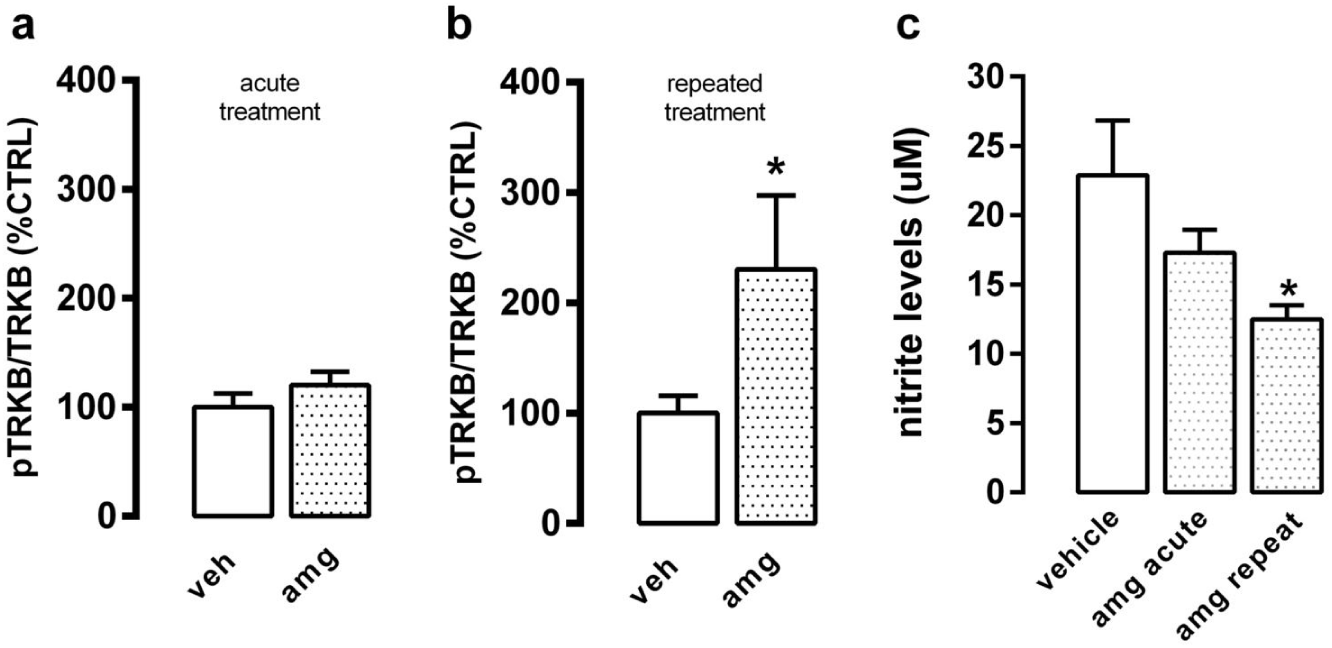
Repeated but not acute treatment with aminoguanidine increases phospho TRKB and decreases nitrite levels in dPAG. Effect of aminoguanidine (15mg/kg, ip) acute (a; n= 5,6) or repeated treatment (b; n= 5/group) on TRKB phosphorylation and nitrite levels (c; n= 11,10,9) in the dPAG. TRKB phosphorylation was normalized by total TRKB levels and expressed as percentage of control (vehicle-treated). Columns represent mean ± SEM; Mann-Whitney (a,b) or Kruskal-Wallis followed by Dunn’s multiple comparison test (c), *p^<^0.05 *vs* vehicle-treated group.

#### Experiment 5: effect of intra-dPAG K252a on AMG panicolytic-like effect

As depicted in figure 3b, the MANOVA showed a significant interaction between the systemic injection of AMG, central administration of K252a and trials [F(1,17)= 5.40, p<0.05] on the escape behavior generated by the ETM. The *post-hoc* test showed that on escape 3, animals of the group vehicle-AMG had longer escape latency when compared to all other groups [F(1,17) = 8.25; Newman-Keuls, p<0.05]. Regarding avoidance (fig. 3a), there was a significant effect of trials [F(1,17)= 25.9, p<0.05], but no effect of systemic AMG [F(1,17)= 0.66, NS] or K252a intra-dPAG [F(1,17)= 2.30, NS]. Also, there was no interaction between the three factors [F(1,17)= 1.08, NS].

**Figure 3.**
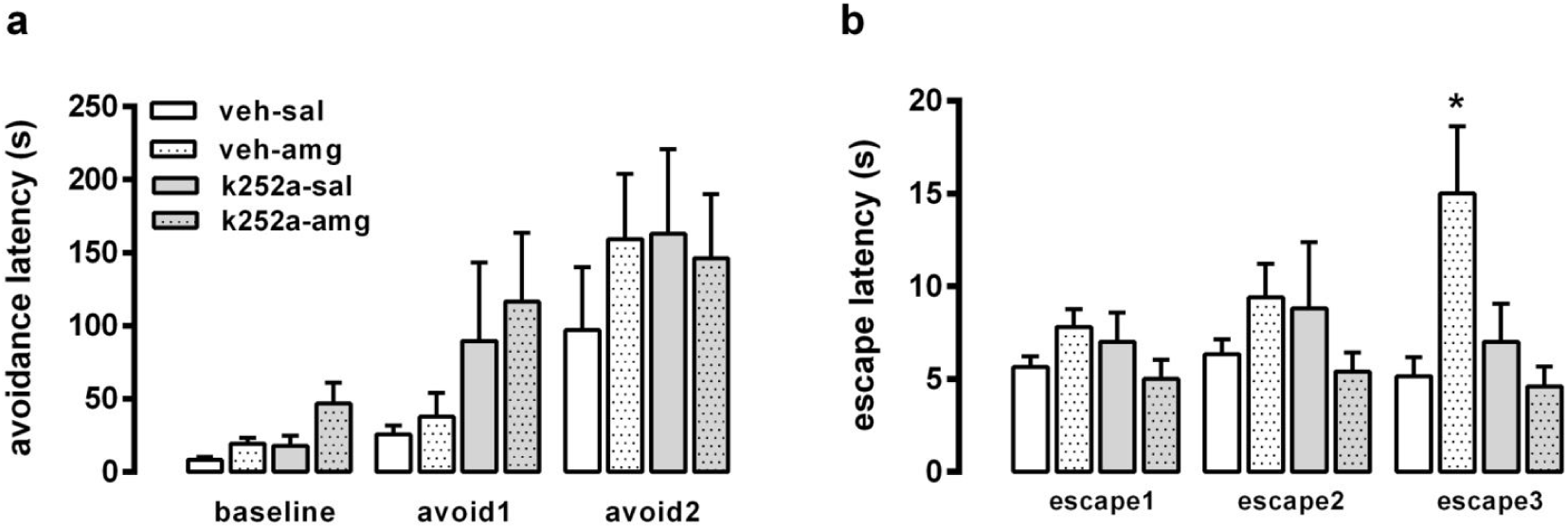
Aminoguanidine-induced panicolytic effect is abolished by the TRKB antagonist K252a. Effect of aminoguanidine (15 mg/kg i.p.) repeated treatment (7 days) and intermittent intra-dPAG injection of K252a (10 pmol/200nl, injected at days 1, 4, 7) on avoidance (a) and escape (b) latencies in ETM (n= 6,5,5,5). Avoidance (a) and escape (b) latencies were expressed as mean ± SEM; two-way repeated measures ANOVA, *p^<^0.05 *vs* vehicle-treated group at the same trial.

Avoidance (time taken to leave the enclosed arm) was measured in 3 consecutive trials (baseline, avoidance I and 2) and expressed in seconds. No snifkant difference was found (repeated two-way ANOV, drug treatment and trials as faciors,

BDNF and TRKB IovIs ar oxrnossod parcontago of th vehicI group. TRB kvols v.,ro not assossod in arng 30 and amg 60 groups. No significant difference was found by one-way ANOVA (BDNF analysis) and t Test (TRKB analysis).

## Discussion

The present study investigated the effect of the non-selective NO synthase inhibitor aminoguanidine (AMG) on the escape response of rats submitted to the ETM and hypoxia tests, and the involvement of TRKB signaling in the AMG-induced effects.

Aminoguanidine is considered a preferential inhibitor of the inducible isoform of NOS (iNOS or NOS2). However, AMG selectivity for iNOS over nNOS is only 4.7 fold (Wolff et al., 1997). Due to the higher expression of nNOS compared to iNOS (Vincent and Kimura, 1992; Onstott et al., 1993), especially in the dPAG, it is most likely that in our experimental conditions AMG is acting mainly upon the neuronal isoform. To our knowledge, there is no significant levels of iNOS in the PAG area, except under stressful conditions such as chronic pain (Mor et al., 2011). Supporting the idea that AMG is inhibiting nNOS in dPAG, we observed decreased levels of nitrite, a direct byproduct of NO production, in the dPAG after AMG repeated treatment, which matched the behavioral effects of repeatedly administered AMG.

AMG inhibited escape response both in the hypoxia test and in ETM. In the hypoxia test, repeated treatment with three doses of AMG decreased escape expression, but this effect reached statistical significance with the lowest dose. Hypoxia test is based on the idea that panic-attacks are a result of a misinterpreted sensation of suffocation, as proposed by Donald Klein (Klein, 1993). In a recent study, Spiacci and colleagues, described the role of the dPAG area in mediating escape responses (jumps) induced by hypoxia exposure. The authors also showed that these responses are attenuated by high potency benzodiazepines (alprazolam) or repeated treatment with fluoxetine (Spiacci et al., 2015) and, in the present study, repeated treatment with AMG. In the ETM, repeated but not acute treatment with AMG 15 or 60 mg/kg induced a clear anti-escape effect that was not observed with 30 mg/kg. The reason for the inefficacy of this intermediate dose is not clear, but similar U- or bell-shaped curves for the effect of NOS inhibitors are not uncommon (Calixto et al., 2008; Lisboa et al., 2013). The frequency of such ‘U-shaped’ and the hormetic model were revised by Calabrese and Baldwin (Calabrese and Baldwin, 2003a, 2003b), and appears to be more common than acknowledged. However, little is known about the molecular mechanisms behind these observations (Rozman and Doull, 2003). It has been proposed that under a challenge (for example, drug treatment) several modifications would be triggered in the nervous system in order to counteract the challenge. However, if the given challenge is too mild (eg. drug treatment with low doses), the counteracting response would not be triggered thus allowing the drug effectiveness. On the other hand, if the challenge is too harsh (eg. higher doses) it will surpass the counteracting response, allowing the drug effectiveness once again. In this scenario, an intermediate dose (enough to trigger the counteracting response but not to surpass it) would result in the inefficacy of the treatment.

Another possibility to explain the lack of effect of the intermediary dose of AMG in LTE escape response is that AMG systemically delivered would differentially affect distinct brains areas, depending on the dose used. Despite the nitrite data implicates NOS inhibition in dPAG, it is not possible to rule out AMG-induced NO inhibition in other brain areas, specially with higher doses. This possibility should be investigated in further studies.

Similar to what we observed with AMG, several clinically used antidepressive/panicolytic drugs, such as fluoxetine, imipramine and escitalopram, inhibit escape expression in the ETM and some of them (e.g imipramine and escitalopram) also impairs inhibitory avoidance acquisition, indicating an anxiolytic effect (Pinheiro et al., 2007). Under the experimental conditions of the present study, however, neither the acute or repeated treatment with AMG were effective in changing this behavioral parameter in the ETM.

Prior studies have investigated the acute effect of NOS inhibitors in the ETM, but as far as we know, there are no other studies investigating the effect of repeated treatment with NOS inhibitors in the ETM. In agreement with our data, no panicolytic-like effect was observed on escape behavior after a single injection of NOS inhibitor L-NAME, either i.p. or intra-dlPAG. However, an anxiolytic-like effect on avoidance response was observed by both i.p. or intra-dlPAG injections (Calixto et al., 2001, 2008). The exact reason for this discrepancy is not fully understood but it might be related to the differences between AMG and L-NAME selectivity upon the NOS isoforms. L-NAME systemically delivered induced a significant increase in the blood pressure which indicates an effect upon endothelial NOS isoform. In fact, as described by Boer and colleagues (Boer et al., 2000), AMG is 6-times more selective to iNOS than eNOS and 5-times for nNOS than eNOS. In the same study the authors observed that L-NAME exhibit no selectivity between nNOS/eNOS isoforms.

Behavioral changes induced by NOS inhibitors have been described after acute and repeated administration, depending on the animal model. For example, NOS inhibitors induce antidepressant-like effect in the forced swimming test after a single injection in mice (Harkin et al., 1999) while repeated treatment is required to observe similar effects in the chronic mild stress or learned-helplessness models (Yazir et al., 2012; Stanquini et al., 2017). Similar profile has been described for classical antidepressant drugs and there is evidence that different molecular mechanisms might be engaged in the effects induced by acute and repeated treatment (Medrihan et al., 2017). In the ETM, it has been consistently found that repeated treatment is required for the anxiolytic and panicolytic effect induced by classical antidepressant drugs, single injection was found ineffective for all the antidepressant drugs tested (Teixeira et al., 2000; Poltronieri et al., 2003; Pinheiro et al., 2007). It has been proposed that the long-term effect of classical antidepressants involves complex neuroplastic alterations in the central nervous system mediated by BDNF-TRKB signaling (Karpova et al., 2011). We hypothesize that regulation of neuronal plasticity might be required for the long-term antidepressant- and anxiolytic/panicolytic-like effects of NOS inhibitors as well, which would explain the lack of effect with acute treatment.

In the proposed scenario, the panicolytic effect of AMG in the ETM could involve the activation of TRKB receptors in the dPAG. Recently, our group described a role for BDNF signaling in the dPAG in panicolytic-like effects caused by antidepressants (Casarotto et al., 2015). Several antidepressant drugs, with anxiolytic and panicolytic properties, such as fluoxetine, are able to increase the levels of pTRKB following acute or short-term treatment regimen in the hippocampus and prefrontal cortex [for review see (Castrén, 2014)]. We observed that pTRKB and BDNF levels were found elevated in the dPAG of imipramine-, but not fluoxetine-treated animals, after 3 days of drug administration. The differential effect between these compounds on pTRKB levels, were also reflected on behavioral experiments. *i.e.* imipramine, but not fluoxetine, exerted panicolytic-like effect in the ETM following this short-term regimen. Both drugs increased pTRKB levels in the dPAG after 21 days of treatment (Casarotto et al., 2015), and are equally effective on ETM at this stage (Pinheiro et al., 2007). The microinjection of BDNF into dPAG increased the escape latency in the ETM (Casarotto et al., 2015) without causing any changes in the avoidance, and this neurotrophin also inhibited the escape response induced by electrical stimulation of dPAG (Casarotto et al., 2010).

Prior studies reported increase in total BDNF levels in hippocampus *in vivo* following repeated treatment with NOS inhibitors (Stanquini et al., 2017). However, we did not found any change in BDNF levels in dPAG after AMG treatment. Despite that, repeated treatment with AMG increased pTRKB in dPAG. The ability of acute and repeated AMG treatment in upregulating pTRK in the dPAG was positively associated with its effectiveness upon the behavioral response observed in the ETM. Moreover, the blockade of AMG-induced TRKB signaling, in dPAG, by k252a blocked also the behavioral effects of AMG on escape latency. Similarly, only this parameter of the ETM was reported to be affected by systemic imipramine or locally injected BDNF, and it was also blocked by intra-dPAG administration of k252a (Casarotto et al., 2015).

Taken together our data suggests that AMG exerts its panicolytic-like effect through an increase in pTRKB levels in the dPAG. It is still unclear how NO could modulate the activation of BDNF receptor but it is plausible to consider that NO regulates local BDNF release in dPAG without changing this neurotrophin levels. Accordingly, Canossa and co-workers (Canossa et al., 2002) showed that NOS inhibition increased BDNF release in cell culture of hippocampal neurons. Another intriguing possibility is a direct modulation of TRKB function by NO, through nitration and/or nitrosylation. Using an algorithm to analyze putative sites for nitration or nitrosylation [developed by (Xue et al., 2010; Liu et al., 2011)] in TRK receptors, we reported previously that TRKB displays sites with high probability for nitration of tyrosine residues (Biojone et al., 2015). Both mouse and rat mature TRKB receptors are potential targets for nitration at Y90, Y342 and Y816 residues. Interestingly, the Y816 residue is also a well-known target of phosphorylation following TRKB activation either by BDNF or antidepressant drugs, which recruits PLC-gamma pathway (Huang and Reichardt, 2001). Functionally, there is no evidence describing changes in nitrated TRKB responses. However, the phosphorylation of TRKA receptors by NGF prevents its nitration by peroxynitrite, suggesting a competitive functional interaction between these two modifications (Spear et al., 2002). Thus, inhibiting the production of NO, putatively reducing the probability of TRKB nitration, could facilitated BDNF effect on TRKB in the dPAG, leading to the observed panicolytic-like effect. This is an interesting hypothesis to explain nitric oxide effects on TRKB phosphorylation, which remains to be explored in future investigation.

## Acknowledgements

The authors thank to Flávia Salata, José C. Aguiar and Eleni T. Gomes for their technical assistance

